# Identification of putative G-quadruplex forming sequences in three manatee papillomaviruses

**DOI:** 10.1101/138602

**Authors:** Maryam Zahin, William L. Dean, Shin-je Ghim, Joongho Joh, Robert D. Gray, Sujita Khanal, Gregory D. Bossart, Antonio A. Mignucci-Giannoni, Eric C. Rouchka, Alfred B. Jenson, Jonathan B. Chaires, Julia H. Chariker

## Abstract

The Florida manatee (*Trichechus manatus latirotris*) is considered a threatened aquatic mammal in United States coastal waters. Over the past decade, the appearance of papillomavirus-induced lesions and viral papillomatosis in manatees has been a concern for those involved in the management and rehabilitation of this species. To date, three manatee papillomaviruses (PVs) have been identified in Florida manatees, one forming cutaneous lesions (TmPV1) and two forming genital lesions (TmPV3 and TmPV4). In this study, we identified DNA sequences with the potential to form G-quadruplex structures in all three PVs. G-quadruplex structures (G4) are guanine-rich nucleic acid sequences capable of forming secondary structures in DNA and RNA. In humans, G4 are known to regulate molecular processes such as transcription and translation. Although G4 have been identified in several viral genomes, including human PVs, no attempt has been made to identify G4 in animal PVs. We found that sequences capable of forming G4 were present on both DNA strands and across coding and non-coding regions on all PVs. The vast majority of the identified sequences would allow the formation of non-canonical structures with only two G-tetrads. The formation of one such structure was supported through biophysical analysis. Computational analysis demonstrated enrichment of G4 sequences on the reverse strand in the E2/E4 region on all manatee PVs and on the forward strand in the E2/E4 region on one genital PV. Several G4 sequences occurred at similar regional locations on all PVs, most notably on the reverse strand in the E2 region. In other cases, G4 were identified at similar regional locations only on PVs forming genital lesions. On all PVs, G4 sequences were located near putative E2 binding sites in the non-coding region. Together, these findings suggest that G4 are likely regulatory elements in manatee PVs.

**Author summary:** G-quadruplex structures (G4) are found in the DNA and RNA of many species and are known to regulate the expression of genes and the synthesis of proteins, among other important molecular processes. Recently, these structures have been identified in several viruses, including the human papillomavirus (PV). As regulatory structures, G4 are of great interest to researchers as drug targets for viral control. In this paper, we identify the first G4 sequences in three PVs infecting a non-human animal, the Florida manatee. Through computational and biophysical analysis, we find that a greater variety of sequence patterns may underlie the formation of these structures than previously identified. The sequences are found in all protein coding regions of the virus and near sites for viral replication in non-coding regions. Furthermore, the distribution of these sequences across the PV genomes supports the notion that sequences are conserved across PV types, suggesting they are under selective pressure. This paper extends previous research on G4 in human PVs with additional evidence for their role as regulators. The G4 sequences we identified also provide potential regulatory targets for researchers interested in controlling this virus in the Florida manatee, a threatened aquatic mammal.

## Introduction

G-quadruplex structures (G4) are four-stranded, inter- and intramolecular structures formed from guanine-rich DNA and RNA sequences. The sequences fold to form stacks of G-tetrads, planar structures composed of four guanine bases held together by Hoogsteen hydrogen bonds, Fig 1 (1). The stacked G-tetrads are connected by loops which vary in size and sequence composition, affecting the stability of the structure.

**Fig 1.**
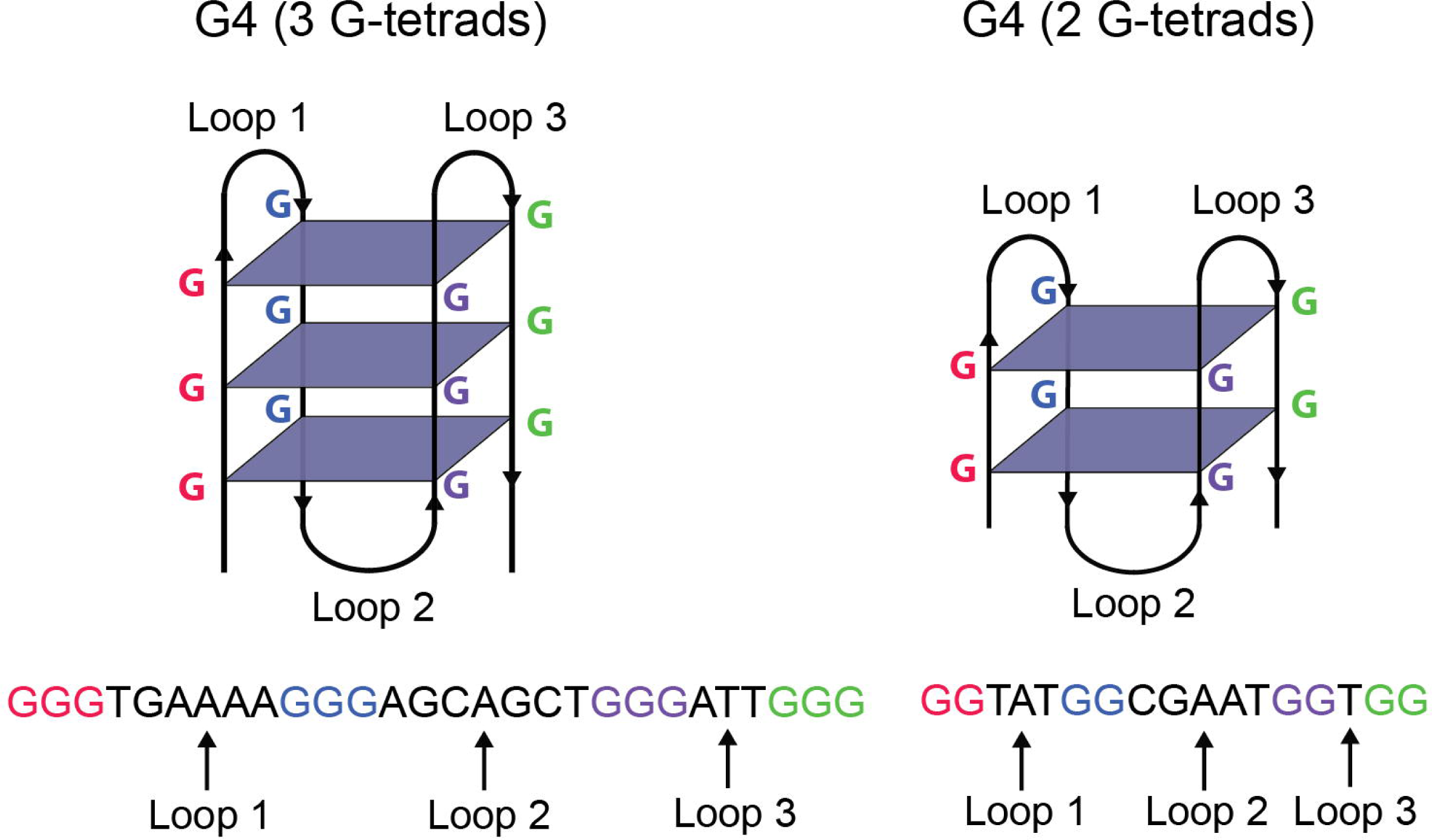
Secondary intramolecular G4 structures (top) and corresponding DNA sequences (bottom) with varying numbers of G-tetrads. This figure has been modified from (54).

G4 are known to be involved in a series of key biological functions. In humans, G4 are found in telomeric repeats and serve to prevent degradation and genomic instability (2). Their formation in this region is also known to decrease telomerase activity which is selectively expressed in a vast majority of cancers (3). G4 located in the promoter region of genes act as transcriptional regulators (4) while those found in intronic and exonic regions play a role in alternative splicing (5, 6). In RNA, G4 identified in 3’ and 5’ untranslated regions are known to regulate protein synthesis (7, 8).

G4 function as regulators through at least a couple of different mechanisms (1). For example, G4 formation can inhibit transcription by blocking the activity of RNA polymerase. Alternatively, G4 can bind with other regulatory elements that either activate or repress transcription. A variety of different proteins are now known to bind with G4 in DNA and RNA (9). In RNA, this includes proteins involved in splicing as well as protein synthesis (10).

A regulatory role for G4 in prokaryotic cells has been well-established (11). This has led to an interest in examining the role G4 may play in organisms such as viruses (12, 13). Although one might presume that viruses have evolved analogous regulatory mechanisms, research on the role of G4 in viral genomes has been limited to human immunodeficiency virus 1 (HIV-1) (14), human papillomaviruses (HPV) (15), and Epstein-Barr virus (EBV) (16). In HIV-1, ligand stabilization of a G4 located in the *nef* gene reduced gene expression and repressed HIV-1 infectivity in an antiviral assay (8). In EBV, destabilization of a G4, located in virally encoded mRNA, reduced translation (11). Both findings are in line with research on the regulatory role of G4 in prokaryotes. In HPV, DNA sequences capable of forming G4 have been identified in four regions (NCR, L2, E1, and E4) of eight HPV types, and their ability to form in the laboratory has been established. However, it remains unclear how they might affect PV replication and transcription (10, 15).

PVs cause a number of benign and malignant tumors in humans and animals. There are approximately 100 human PV types and at least 112 non-human PV types found in 54 different species (17). The specific location of tumor formation (cutaneous, oral, genital, or anal) depends on the type of PV. The existence of G4 in non-human animals would provide further support for their potential functional relevance in PVs and would also provide a valuable comparison to the pattern of distribution seen in human PVs. In the current paper, we identify and characterize G4 sequences in three PVs infecting the Florida manatee.

The study of G4 in manatee PVs has ecological as well as biological significance. The Florida manatee is an aquatic mammal living in the coastal waters of Florida that has been classified as an endangered species since 1967. Its population declined for a variety of reasons, not the least of which was that its gentle, slow-moving nature made it vulnerable to injury from boat propellers. Efforts at restoring the population have been successful to the extent that the species was downlisted to threatened status in 2016. However, in the midst of these efforts, animals undergoing rehabilitation frequently showed signs of high sensitivity to environmental stress, one sign being the development of cutaneous or mucosotropic genital papillomatous lesions. Some animals, both in captivity and in the wild, showed antibody titers indicating the presence or exposure to *Trichechus manatus* PV 1 (TmPV1), a virus first characterized in our laboratory (18). More recently, genital lesions appeared in a single Florida manatee used as a surrogate animal for manatee rehabilitation at the Puerto Rico Center (PRMCC), and DNA sequencing, also performed in our laboratory, indicated the presence of two new PVs, *Trichechus manatus* PV 3 (TmPV3) an*d* 4 (TmPV4) (19, 20). These are the first known genital mucosotropic PVs in a manatee, presenting a potential health threat to this species should the virus spread in wild populations, if not already present there.

Similar to HPVs, manatee PV genomes are comprised of double stranded DNA, approximately 8 Kb in length that encodes a maximum of seven genes. Five genes encode non-structural or early proteins E1, E2, E4, E6 and E7, and two encode structural or late proteins L1 and L2, with all coding regions located on the forward DNA strand. A non-coding region holds the origin of replication and at least a couple of promotor sites. Much of what is known about the function of these sites comes from molecular biological research on human PVs (21). E1 and E2 proteins form a complex that initiates viral replication at the origin, resulting in amplification of the virus. E2 also functions as a negative regulator of E6 and E7, two coding regions that stimulate cell growth and function as oncogenes in human PVs. The late proteins L1 and L2 code for the major and minor capsid proteins encapsidating viral DNA with E4 having a possible role in facilitating virion release.

During our initial sequencing of TmPV4, a glycine rich GGA repeat sequence identified in the E2 region created an obstacle to sequencing due to the formation of a secondary structure, necessitating the use of power-read sequencing analysis to complete the genome (19, 20). We reasoned this was likely to be a G4, given that a GGA repeat would be a sequence pattern likely to form one of these structures. In fact, in 2001 Matsugami and colleagues reported the folding of a GGA repeat into an intramolecular parallel G-quadruplex in the laboratory (22). This is significant in that a G4 in the E2 region could play a vital role in altering the regulatory functions of the virus due to the role E2 has in regulating the expression of oncoproteins E6 and E7. Moreover, integration of E2 into a human cervical host cell chromosome is considered by epidemiologists to be an important event leading to the development of cervical cancer. From a functional standpoint, blockage of E2 transcription by integration could potentially be equivalent to blockage of E2 by a G-quadruplex structure.

In this paper, we identify sequences with the potential to form G4 on both DNA strands in each coding and non-coding region on all three manatee PV genomes. In contrast to the findings for G4 in HPV, we find that the majority of sequences identified were capable of forming G4 structures with two rather than three G-tetrads, and we provide laboratory support for the formation of a secondary structure from one such sequence. We find several G4 in similar locations on all three PVs as well as several G4 in similar locations unique to the two PVs forming genital lesions. G4 were also located near putative E2 binding sites in non-coding regions on all PVs. Although G4 were found in all coding and non-coding regions, G4 were significantly enriched in the E2/E4 region on all three genomes, suggesting that G4 are evolutionarily preserved in this region.

## Results

### G4 sequence distribution

The number of putative G4 sequences, broken down by DNA strand, genomic region, and TmPV genome, is displayed in Table 1. As described in the Materials and methods section, longer sequences supported the development of more than one G4 at a time. As a result, in several regions, the number of G4 possible was slightly higher than the number of sequences identified, and these values are displayed alongside the number of G4 sequences in Table 1. TmPV4 had the highest number of sequences identified, with 20 on the forward DNA strand and 17 on the reverse strand. Somewhat fewer sequences were identified on TmPV3 (13 forward, 11 reverse) and TmPV1 (14 forward, 15 reverse).

**Table 1.** The number of putative G4 sequences identified and the number of structures possible across different regions on forward and reverse DNA strands for each TmPV genome.

| Region | Forward DNA Strand<br>Number Sequences (Number<br>Possible Structures) |  |  | Reverse DNA Strand<br>Number Sequences (Number<br>Possible Structures) |  |  |
| --- | --- | --- | --- | --- | --- | --- |
|  | TmPV1 | TmPV3 | TmPV4 | TmPV1 | TmPV3 | TmPV4 |
| E6 | - | 1 (1) | - | - | - | - |
| E7 | 2 (2) | 1 (1) | - | - | - | - |
| E1 | 5 (5) | 3 (3) | 4 (5) | - | - | - |
| E2 | 1 (1) | 1 (1) | 2 (2) | 1 (1) | - | - |
| E1/E2 | 1 (1) | 1 (1) | 1 (1) | - | - | - |
| E2/E4 | 1 (1) | - | 7 (12) | 3 (5) | 3 (6) | 6 (9) |
| Total<br>Early Region | 10 (10) | 7 (7) | 14 (20) | 4 (6) | 3 (6) | 6 (9) |
| L2 | 2 (4) | 2 (4) | 3 (3) | 5 (6) | 5 (6) | 7 (7) |
| L1 | 2 (2) | 3 (3) | 2 (2) | 4 (5) | 3 (3) | 3 (3) |
| Total<br>Late Region | 4 (6) | 5 (7) | 5 (5) | 9 (11) | 8 (9) | 10 (10) |
| NCR | - | 1 (1) | 1 (3) | 2 (2) | - | 1 (1) |
| Total Genome | 14 (16) | 13 (15) | 20 (28) | 15 (19) | 11 (15) | 17 (20) |

All identified G4 sequences, with one exception, are capable of forming G4 with only two G-tetrads. The exception to this pattern was found in the L2 region of TmPV1 where a sequence capable of forming a three G-tetrad structure on the forward DNA strand was identified. This sequence was embedded in a much longer sequence also capable of forming a two G-tetrad structure. The individual sequences along with sequence locations and sequence descriptors for putative G4 identified in each TmPV genome are available in S1, S2, and S3 Tables.

### G4 enriched in E2/E4 region on all TmPVs

For all TmPVs, the number of nucleotides covered by G4 sequences was greater than expected in the E2/E4 region when compared to a random distribution of G4 across the genome. This occurred primarily on the reverse DNA strand (E2: TmPV1, *p* = 0.05; TmPV3, *p* = 0.053; TmPV4, *p* = 0.015; E4: TmPV1, *p* = 0.005; TmPV3, *p* = 0.008; TmPV4, *p* = 0.001). However, TmPV4 also showed enrichment on the forward DNA strand (E2: *p* = 0.013; E4: *p* = 0.009). The number of observed G4 nucleotides, the number of random simulations with G4 nucleotides greater than or equal to the observed G4 nucleotides, and the associated significance values are available in S4 Table for each DNA strand in each genomic region on each TmPV.

### Co-occurring G4 locations across TmPVs

To identify similar patterns in the distribution of G4 sequences across the three manatee PV genomes, the coding/non-coding regions were aligned at their starting locations, and G4 sequences with one or more nucleotides at the same distance from the beginning of the region were identified as occurring at the same location within a region. There were several regions with G4 sequences in the same location on all genomes. In Fig 2, on the forward DNA strand (left), G4 sequences were found in the same location in E2, L2, and L1. On the reverse DNA strand (Fig 2, right), G4 sequences were found in the same location in E2 and L1. There were also regions with G4 sequences occurring in the same location on TmPV3 and TmPV4 but not TmPV1. On the forward strand this pattern was found in E1 and NCR, and on the reverse strand, this pattern was found in E4.

**Fig 2.**
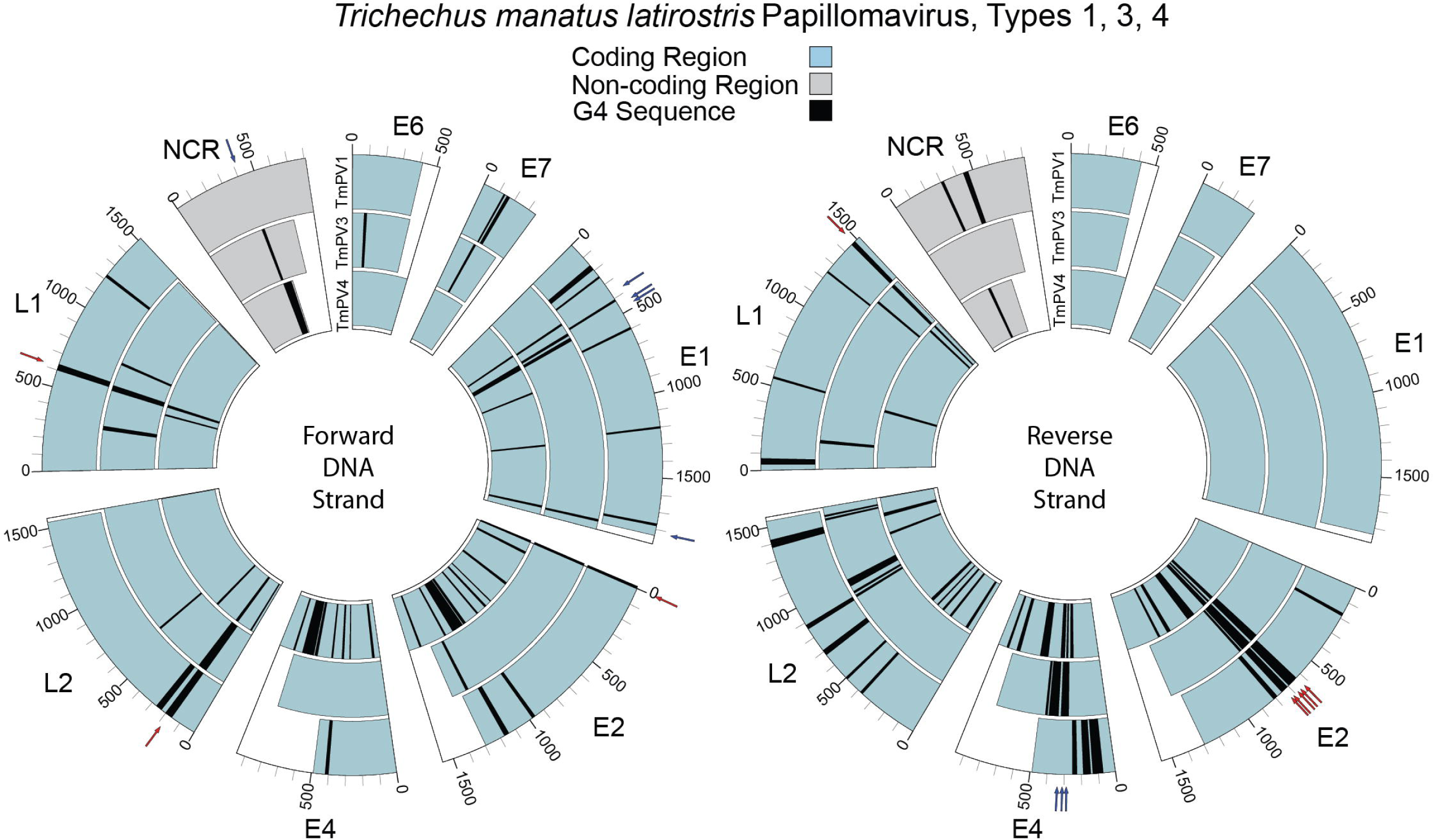
Coding and non-coding regions aligned across TmPV1 (outer), TmPV3 (middle), and TmPV4 (inner) to illustrate G4 sequences occurring at the same location within regions on the forward (left) and reverse (right) DNA strands. Blue arrows indicate G4 sequences at the same location on TmPV3 and TmPV4. Red arrows indicate G4 sequences at the same location on all three genomes.

### G4 located near putative E2 binding sites

On each PV, G4 sequences were identified in the non-coding region (NCR) where the origin of replication is located. Given the known role of G4 in replication (23, 24), these regions were searched for E2 binding sites to determine whether the G4 sequences might be positioned close to the origin of replication. E2 binding sites were identified using the consensus sequence ACCgNNNNcGGT, allowing some variation in the fourth and ninth nucleotide positions (lowercase g and c) with most variation occurring from nucleotide positions 5 through 8 (25). Table 2 lists all sequences identified with the pattern ACCNNNNNNGGT. Nine of the 21 sequences identified have the more conservative consensus sequence of ACCGNNNNCGGT. The locations of consensus sequences identified in the NCR are displayed in Fig 3 along with the location of G4 in that region. On each PV genome, one or more G4 are located within 100 nt of a putative E2 binding site.

**Fig 3.**
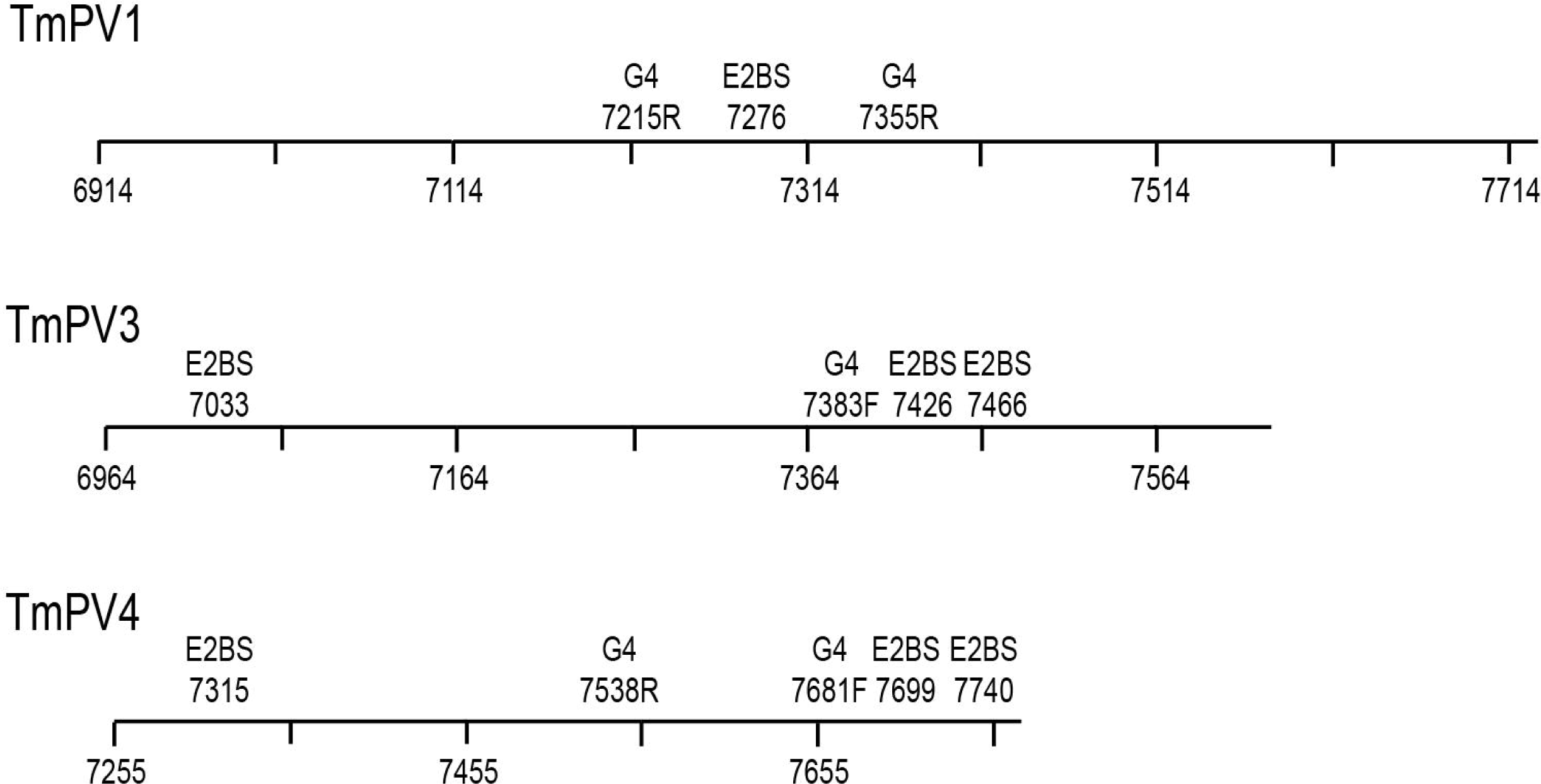
Location of G4 sequences in relation to putative E2 binding sites in the NCR region of each manatee papillomavirus. Only locations where the more conservative sequence ACCGNNNNCGGT is found are displayed.

**Table 2.**
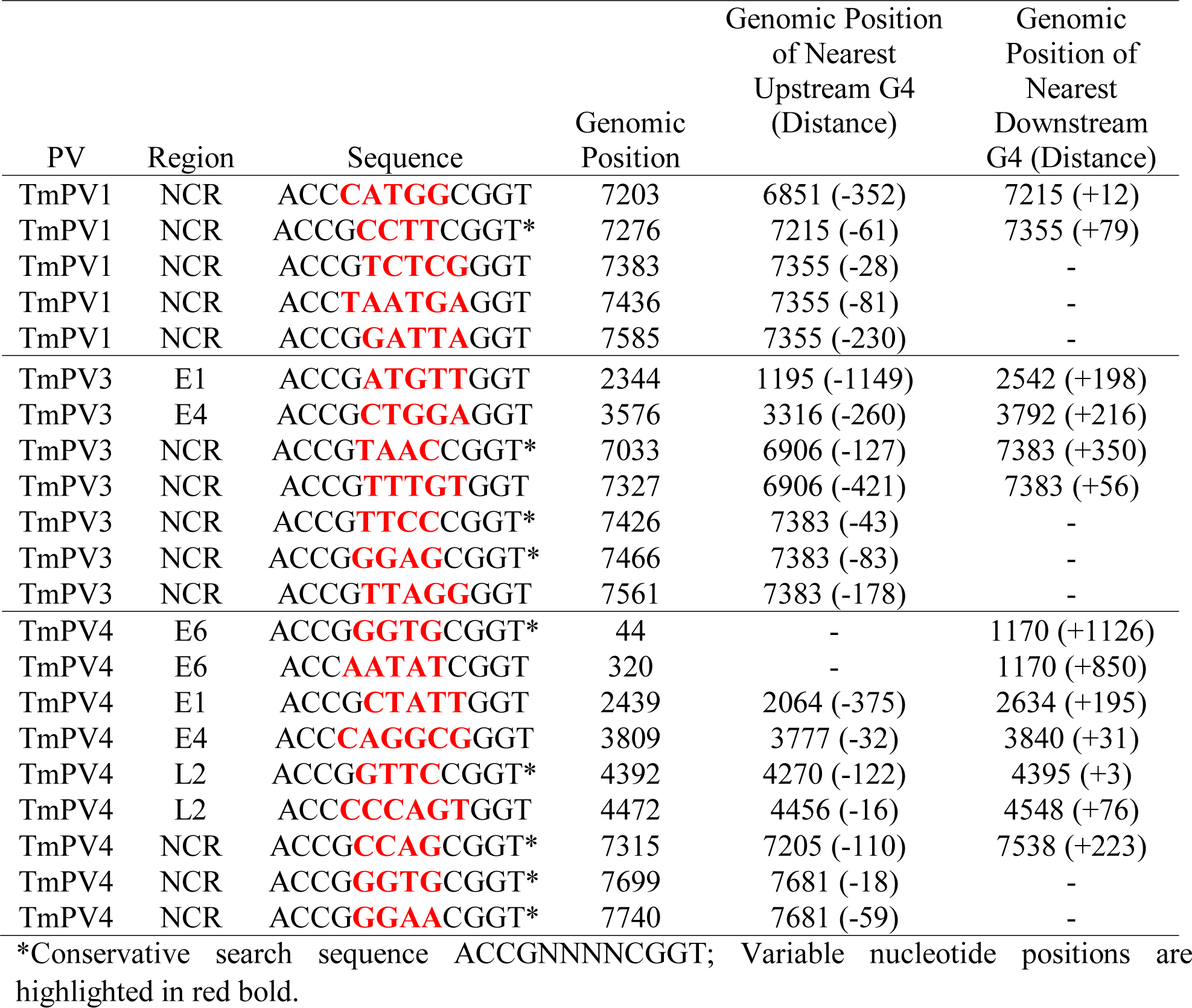
Putative E2 binding site sequences and locations on three manatee PVs along with the location and distance of the nearest putative G4 sequence.

### TmPV4 laboratory analysis

To study the possible formation of G4 secondary structures, analytical ultracentrifugation (AUC) was performed on three oligonucleotides selected from the TmPV4 E2 region, TmPV4-1 (Mw=11,453), TmPV4-2 (Mw=22,045), and TmPV4-3 (Mw=3,511). TmPV4-1 sedimented as a mixture of several molecular species as monomers (total accounted for 90% monomers, Fig 4A). TmPV4-2 was also a mixture of structures, about half monomeric and the rest aggregated (57% monomers, Fig 4B). In contrast, TmPV4-3 sedimented as a 90% dimer (Fig 4C), indicating the possible formation of a two-stranded G-quadruplex structure. The circular dichroism (CD) spectrum is consistent with such a structure as illustrated in Fig 5 and 6.

**Fig 4.**
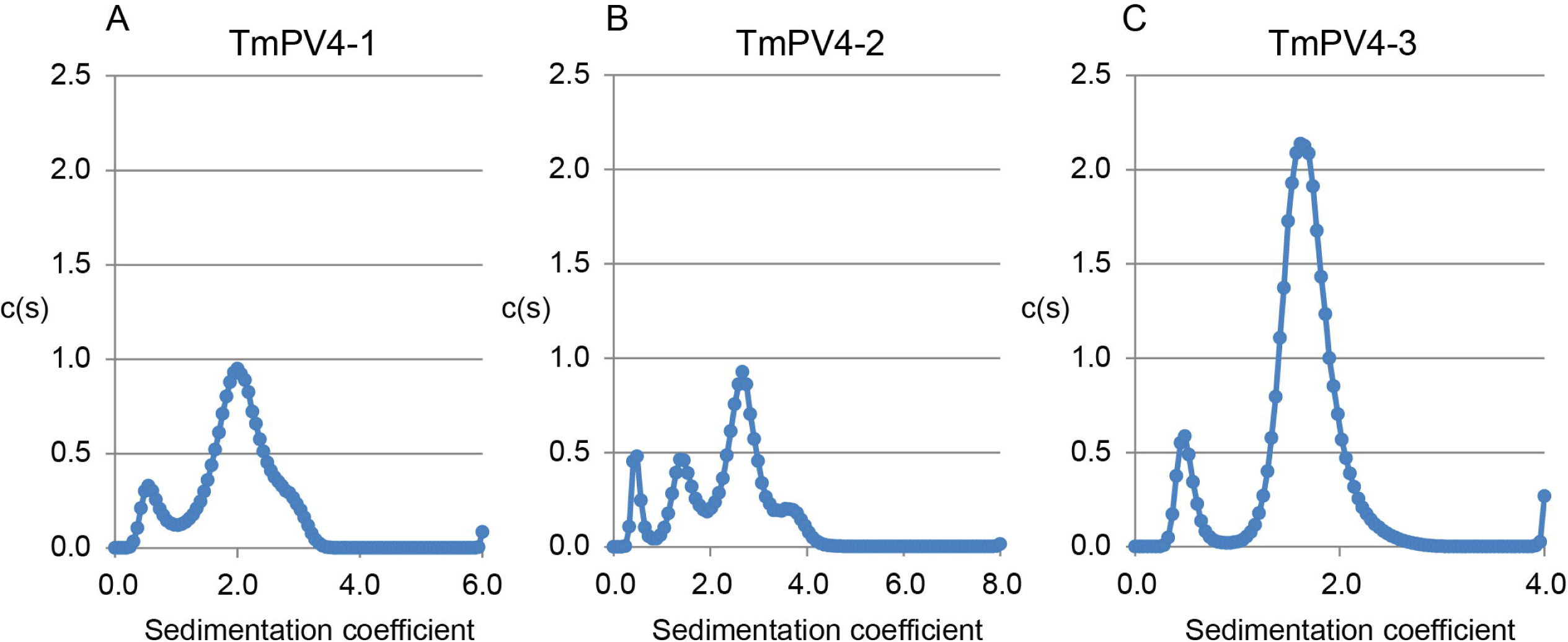
Analytical centrifugation (AUC) of TmPV4 oligos (A, B, and C).

**Fig 5.**
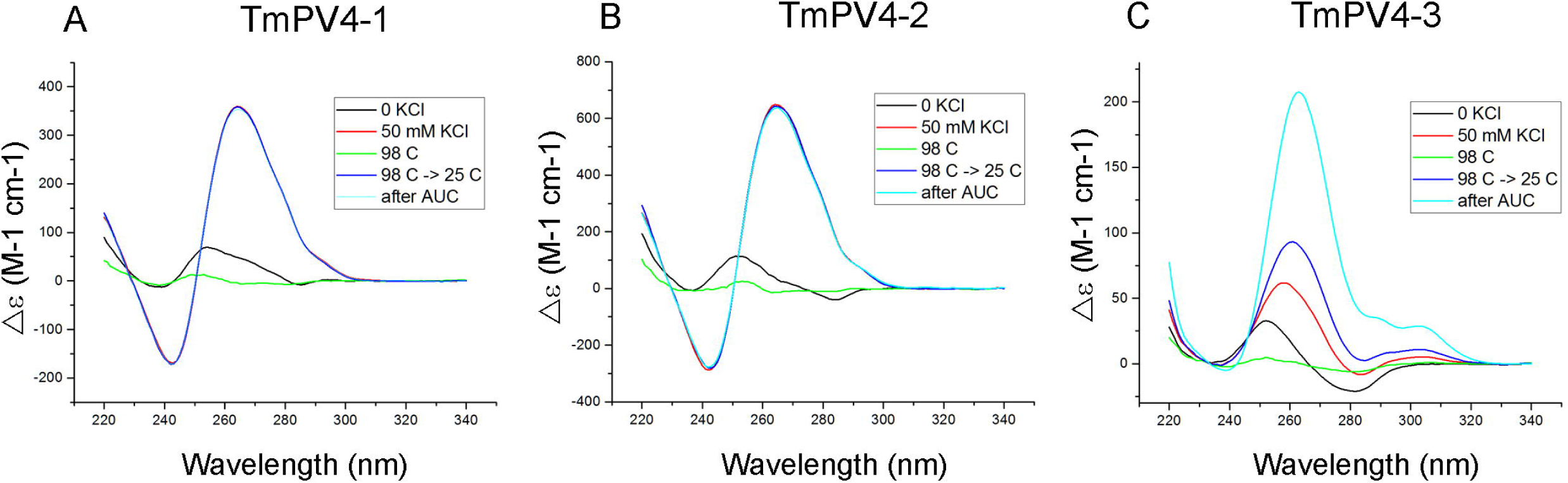
CD-Spectrum analysis of TmPV4 oligos (A, B, and C). Analysis was performed in tBAP/1 mM EDTA/50 mM KCl/pH 7.0.

**Fig 6.**
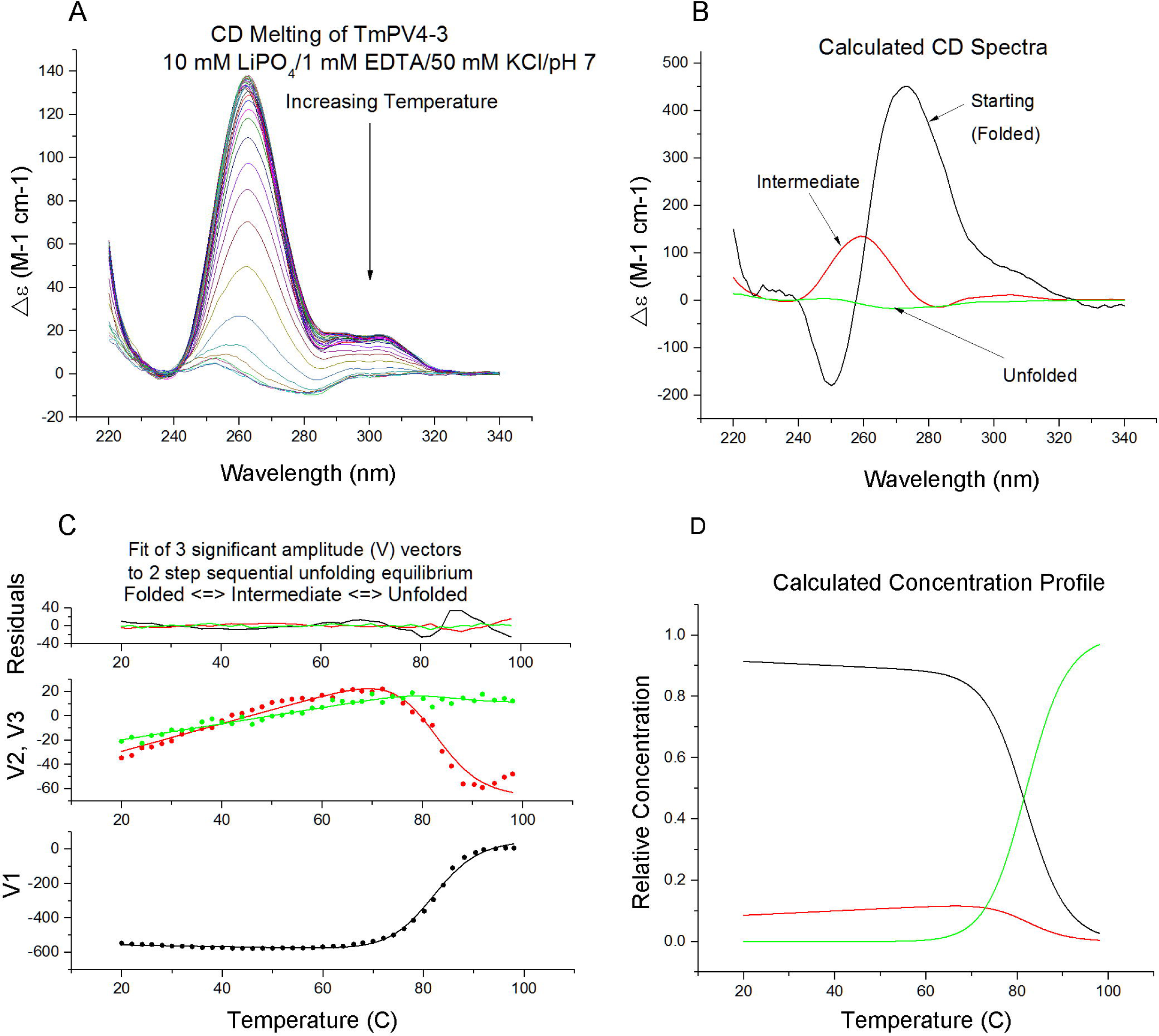
Melting curve (SVD) analysis of temperature-dependent CD spectra of TmPV4-3. Analysis was performed in 10 mM LiPO4/1 mM EDTA/50 mM KCl/pH 7.0. Optimized parameters were ΔH1 = -56.6, Tm1 = 81.42, ΔH2 = -55.05, Tm2 = 72.98, ΔG1 = -9.8, ΔG2 = -8.43, ΔGtotal = -18.23.

## Discussion

The current study represents the first G4 sequences identified in PVs infecting a non-human animal. Prior to this, G4 sequences had been identified solely in PVs infecting humans (15). The G4 sequences identified in the current study consisted, almost exclusively, of sequences capable of forming structures comprised of two G-tetrads, whereas the authors of the recent study of G4 in human PVs searched only for canonical structures comprised of three G-tetrads. As reported in the study, these three G-tetrad sequences were identified in only eight of all human PV types listed in the NCBI Entrez Gene database. However, two G-tetrad sequences are found throughout the human genome (26) and are known to form secondary structures (27). In at least one case, these two-G-tetrad structures have been found to be more stable than three G-tetrad structures (28). In a second case, a similar structure with different loop interactions and capping structures was identified (29), supporting the notion that sequences capable of forming G4 are more variable than once thought. This suggests that the search for G4 in PVs should be extended to include two-tetrad structures.

Our biophysical analysis confirmed that one such sequence in TmPV4, a GGA4 motif located on the forward strand of the E2/E4 region, formed a secondary structure in the laboratory. GGA repeat sequences are common across eukaryotic genomes (30), and the biological relevance of GGA repeats is well-known (31–35). One GGA repeat sequence has been found to form an intramolecular G4 in the laboratory (22, 36). A second GGA repeat sequence located in the *c-myb* promotor forms a G4 that may repress *c-myb* activity through interaction with the *myc*-associated zinc finger protein (37).

In the current study, sequences capable of forming structures with two or more G-tetrads are located more broadly across coding regions than is seen in human PVs when searching for three G-tetrad structures. In the human viruses, three G-tetrad sequences were identified only in the E1 region on the forward DNA strand (15), whereas in manatee PVs we find two G-tetrad structures across all coding regions. Interestingly, the pattern we find on the reverse DNA strand for manatees is somewhat similar to that seen in human PVs; no sequences capable of forming G4 structures are found in E6, E7, or E1. In manatee PVs, all subsequent regions contain two G-tetrad sequences. In human PVs, three G-tetrad sequences are found in E2/E4, L2, and the non-coding region.

The distribution of G4 sequences within coding and non-coding regions across the viruses suggests that G4 may have a variety of regulatory roles. G4 sequences located on the forward DNA strand that are transcribed to mRNA could regulate splicing and translation of early and late genes by binding splicing factors or other related proteins or by serving to block the machinery necessary for each of these processes. In one other virus, Epstein-Barr virus, G4 have been found to inhibit translation of mRNA (16) making this an important avenue to explore in future studies. Similarly, on the reverse DNA strand, the formation of G4 could serve to inhibit transcription through blocking polymerase or binding proteins that enhance or inhibit transcription. In all cases, G4 serve as valuable potential drug targets for viral control.

On all three manatee PVs, G4 were identified at the same location on the forward DNA strand in the L1 and L2 coding regions, perhaps indicating a conserved function for these sequences in the formation of the capsid proteins. New evidence suggests that G4 have a role in immune evasion through antigenic variation and viral silencing (13). In *Neisseria gonorrhoeae*, a G4 structure is essential to variation in the surface protein pilin, allowing the bacteria to evade detection by the host immune system (38). The proposed mechanism involves transcription of an sRNA from the G4 site that base-pairs with the complementary location on the opposite DNA strand, separating the two strands and allowing the formation of the G4. The G4 secondary structures are known to be mutagenic (39), and in HIV-1 and HIV-2 mutated capsids are known to affect the ability of dendritic cells to detect the virus (40).

G4 sequences were also identified in non-coding areas on all three manatee PVs and were located near putative E2 binding sites, indicating a potential role in replication and/or transcription initiation. This would not be surprising given that G4 have been associated with origins of replication in mouse and humans (24, 41). In a study of two vertebrate replicators, G4 were required for initiation of replication and determination of the replication start site (23). On the two manatee PVs producing genital lesions, three putative E2 binding sites were located in a pattern similar to that of genital human PVs (42). One E2 binding site was located on the 5’ end of the non-coding region and two adjacent E2 binding sites were located on the 3’ end separated by 6 and 28 bases. Interestingly, a G4 sequence is located just upstream of the two adjacent 3’ E2 binding sites in an area that should be near the origin of replication. The location of these G4 within the PV promoter region may also be indicative of a role in transcriptional regulation. G4 are found in many promoter sites in humans and are known to regulate transcription (43). In HPV16, two adjacent E2 binding sites in this area were found necessary for negative transcriptional regulation (44).

Interestingly, G4 sequences were enriched on the reverse DNA strand in the E2/E4 regions, suggesting an evolutionary advantage for G4 in this region. This is a significant location given that E2 is a negative regulator of E6/E7, two potential oncogenes. However, the formation of G4 on the template DNA strand would appear to serve simply as a block to the polymerase, inhibiting expression of the early genes, and it is not clear how this would provide an evolutionary advantage to the virus. Alternatively, G4 are known to have a mutagenic effect during replication (45), and G4 sequences in this area may provide some evolutionarily advantageous disruption of the sequence in this region.

## Conclusion

This study provides further confirmation of the existence of G4 in PVs. It extends previous work in human PVs by demonstrating the existence of non-canonical sequences, more broadly located across the genome, that are capable of forming G4 in non-human PVs. Distribution patterns that are indicative of G4 sequence conservation in specific coding regions support the notion that these structures regulate activities similar to those of G4 in other species. As regulatory structures, G4 offer potential drug targets for researchers interested in controlling disease processes (4). Given the threatened status of the Florida manatee and the concerns of scientists working to protect the health of this species, this research also provides an important first step in exploring the biological significance of these structures in this gentle aquatic mammal.

## Materials and methods

### Bioinformatics

The complete genomes for *Trichechus manatus* PVs (TmPV) 1, 3, and 4 were obtained from the NCBI nucleotide division of GenBank (46). Information for accessing individual sequences can be found in Table 3.

**Table 3.** GenBank Sequence Information for TmPV Genomes.

| Accession no. | Description | Size |
| --- | --- | --- |
| NC_006563.1 | TmPV1 | 7722 bp |
| KP205502.1 | TmPV3 | 7622 bp |
| KP205503.2 | TmPV4 | 7771 bp |

Putative G4 forming sequences were identified along each genome using the Quadparser algorithm (47). For each genome, two separate analyses were performed to identify sequences capable of forming unimolecular structures with two and three G-tetrads. In each case, loop lengths were restricted to one to seven bases. The regular expression (G{2,}[ATGC]{1,7}){3,}G{2,} was used to identify structures with two G-tetrads, and the regular expression (G{3,}[ATGC]{1,7}){3,}G{3,} was used to search for structures with three G-tetrads. Quadparser was instructed to search for G4 sequences on the forward DNA strand by searching for runs of guanine bases. However, G4 sequences on the reverse DNA strand were identified by searching for runs of cytosine, guanine’s complement.

Sequences identified by Quadparser generally vary in the number of guanine tracts. A G4 requires four guanine tracts to form. Therefore, sequences with five or more guanine tracks can form G4 at different locations in the sequence, and sequences with eight or more guanine tracts support the development of more than one G4 at a time. To highlight these variations, Quadparser provides a sequence descriptor in the form *x:y:z* (Fig 7), indicating the number of guanine tracts in the sequence (*x*), the number of locations at which a G4 could form (*y*), and the number of G4 possible at a given time within the sequence (*z*).

**Fig 7.**
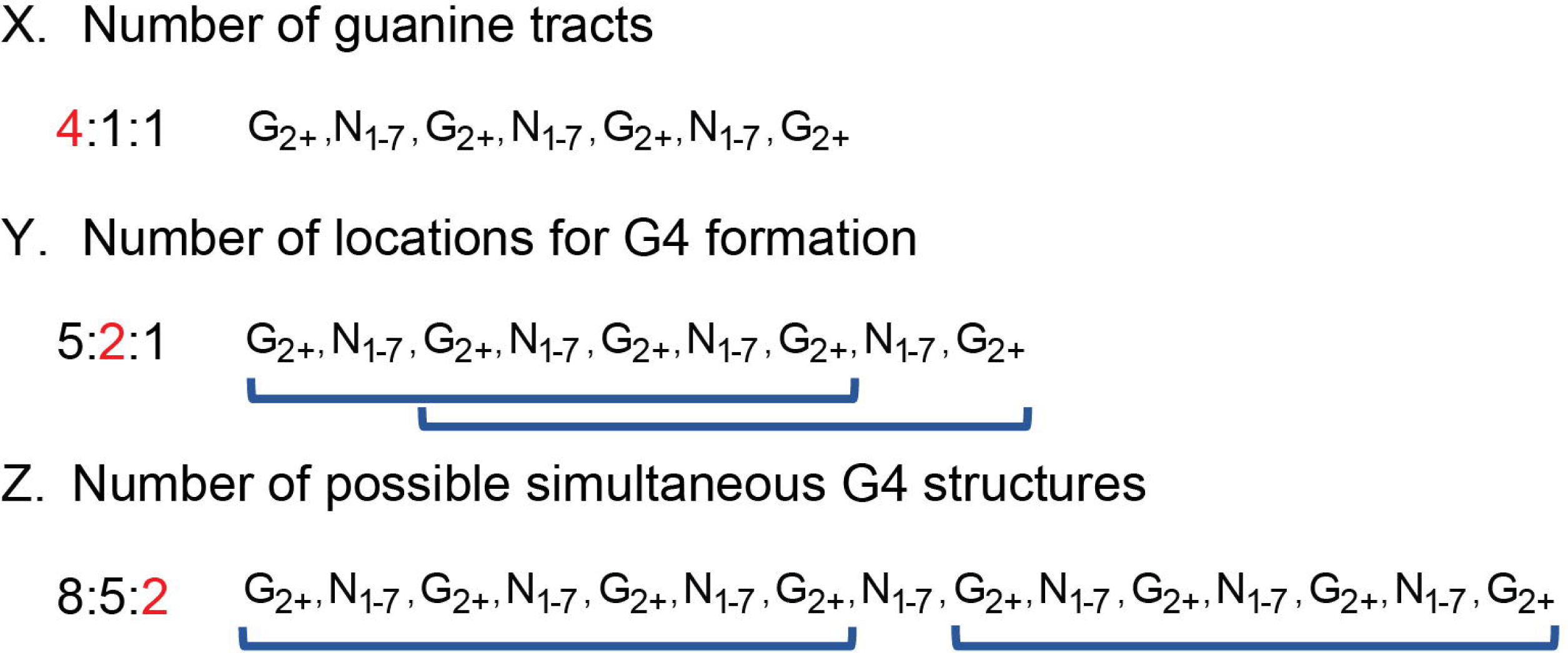
A description of the sequence codes provided by Quadparser for each putative G4 sequence. This figure has been modified from (54).

To establish values for the number of G4 sequences expected in each genomic region at random, Bedtools v2.16.2 (48) was used to randomly distribute the G4 sequences along each PV genome 1,000 times without overlap of the sequences. A Perl program was written to count the number of nucleotides covered by G4 sequence in each coding/non-coding region on each DNA strand for each of the random distributions of G4 sequences and for the observed G4 sequences identified by Quadparser. Significance values for G4 sequence coverage in each region were estimated as *P* = (*r* + 1)/(*n* + 1), where *r* is the number of random simulations in which the coverage of G4 sequences in a region was greater than or equal to the observed coverage of G4 sequences in that region and *n* is the number of random simulations (49, 50). Significance levels were calculated for G4 sequence coverage on each DNA strand in each region of the PV genomes.

Putative E2 binding sites were identified using the consensus sequence 5’-ACCgNNNNcGGT-3’ derived from a comprehensive study of human PVs (25). The consensus sequence includes some variation in the fourth and ninth nucleotide positions with most variation occurring from nucleotide positions 5 through 8. To cast the widest net in searching for E2 binding sites in manatee PVs, positions 4 through 9 were allowed to vary over all nucleotides in the regular expression designed to search for these sequences.

### Oligonucleotides

Oligodeoxynucleotides TmPV4-1, TmPV4-2 and TmPV4-3 (sequences are given in Table 4) were obtained as desalted, lyophilized solids from Eurofins MWG Operon (Huntsville, AL). Each was reconstituted in MQ H20 to give ~1 mM stock solutions based on the manufacturer’s yield. Concentrations were estimated from the absorbance at 260 nm of suitable dilutions into K+-free tBAP buffer (10 mM tetrabutyl ammonium phosphate, 1 mM EDTA, pH 7.0) in conjunction with extinction coefficients supplied by the manufacturer. For NMR experiments, the oligonucleotide was dialyzed vs. 10 mM LiPO4, 50 mM KCl, pH 7.0, prior to measurement.

**Table 4.**
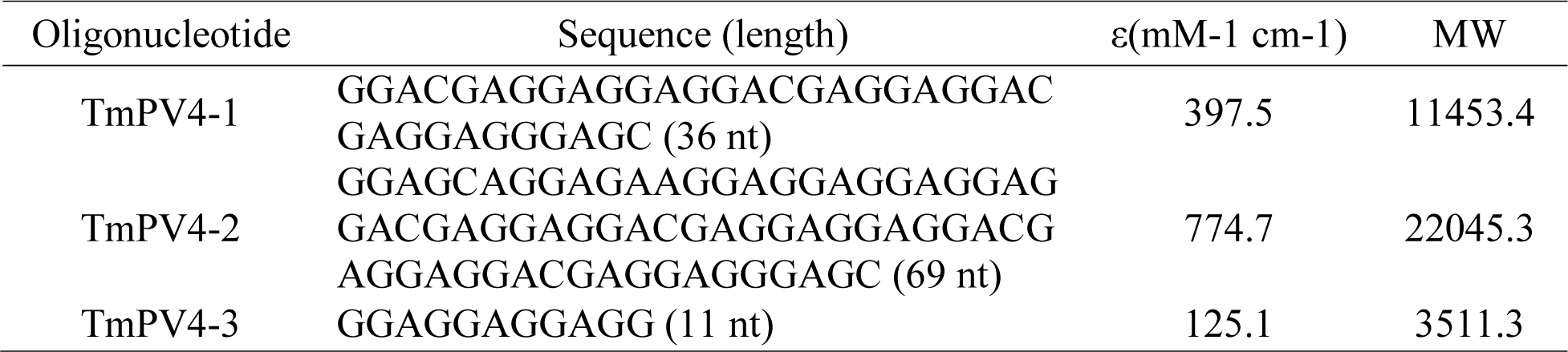
Oligodeoxynucleotides used in this study.

### Analytical ultracentrifugation (AUC) method

Sedimentation velocity experiments were carried out in a Beckman Coulter ProteomeLab XL-A analytical ultracentrifuge (Beckman Coulter Inc., Brea, CA) at 20°C and 50,000 rpm in standard 2 sector cells. Buffer density was determined on a Mettler/Paar Calculating Density Meter DMA 55A at 20.0 °C and viscosity was measured using an Anton Parr AMVn Automated Microviscometer at 20°C. Data were analyzed with the program Sedfit (free software: www.analyticalultracentrifugation.com) using the continuous c(s) distribution model. A value of 0.55 ml/g was used for the DNA oligonucleotides as described (51).

### Circular dichroism spectra

Ultraviolet (UV) and circular dichroism (CD) spectra were measured at 25 °C in a stoppered 1-cm cuvette with a Jasco J-810 spectropolarimeter equipped with a programmable Peltier thermostatted cuvette holder and magnetic stirrer. Instrumental parameters were: 1.0 nm bandwidth, 2 s integration time, 200 nm/min scan rate, four scans averaged. CD data were corrected by subtracting a buffer blank and then normalized using the relationship ε_L_ − ε_R_ = Δε =θ/(32980·c·l), where θ is the observed ellipticity in millidegrees, c is the DNA strand concentration in mol·L^−1^, and l is the path length in cm (52).

### Circular dichroism melts

Thermal denaturation studies of oligonucleotide TmPV4-3 in 10 mM LiPO4, 50 mM KCl, pH 7.0, were carried out essentially as previously described (52). CD spectra were recorded from 320 nm to 220 nm over the temperature range of 4°C to 98 °C at intervals of 1 °C using the instrumental parameters described above. Thermal denaturation was carried out with a temperature ramp of 4 °C/min, ±0.05 °C equilibration tolerance and 60 s delay after equilibration. The resulting temperature/wavelength data matrices were analyzed by singular value decomposition (SVD) as described (53) to obtain melting temperatures (Tm) and enthalpy values (ΔH). Two melts of the same sample were carried out on subsequent days to assess reversibility of the thermal denaturation process.

## Acknowledgements

The original sampling of TmPV3 and TmPV4 was carried out under USFWS permit #MA231088.

## Supporting information captions

**S1 Table. Sequences, locations, and descriptors for putative G4 identified on TmPV1.** Note that all sequences are identified on a reference genome. Thus, G4 sequences on the reverse DNA strand are identified by searching for C-tracts.

**S2 Table. Sequences, locations, and descriptors for putative G4 identified on TmPV3.** Note that all sequences are identified on a reference genome. Thus, G4 sequences on the reverse DNA strand are identified by searching for C-tracts.

**S3 Table. Sequences, locations, and descriptors for putative G4 identified on TmPV4.** Note that all sequences are identified on a reference genome. Thus, G4 sequences on the reverse DNA strand are identified by searching for C-tracts.

**S4 Table. The number of observed G4 nucleotides (NT), the number of random simulations with G4 nucleotides greater than or equal to the observed G4 nucleotides, and the associated significance values for each DNA strand in each genomic region on each TmPV.**

